# *Rectus Femoris* Muscle Elasticity and Stiffness Correlates with Maximal Oxygen Consumption in Triathletes

**DOI:** 10.1101/2023.01.20.524872

**Authors:** Georg Gavronski, Ain Reimets, Jaak Talts, Indrek Koovit, Tõnis Mandel, Ragnar Viir, Eero Vasar, Alar Veraksitš

## Abstract

VO2max is considered single best indicator of cardiovascular fitness and aerobic endurance. We analyzed retrospectively, are there any relationships between muscle parameters and oxygen consumption in a study where the myoton equipment was used to establish muscle biomechanical properties, such as elasticity, stiffness, and tension (measured as oscillation frequency) in triathletes. Eight muscles were studied in 14 male triathletes over three years. Relaxed and contracted states of muscles were measured. VO2max was recorded in these athletes up to four times during this period. Average values were calculated for each athlete and High (max 71.8–min 62.3 ml/kg/min) and Low (59.1–51.3) oxygen consumption groups were formed. Higher oxygen consumption correlated significantly (r=−0.58; p=0.029) with improved elasticity (represented by smaller decrement values) of the *rectus femoris* muscle in a contracted state. Also, in the High VO2max group, this muscle (in a relaxed state) was significantly more elastic and stiffer at the same time compared to the Low group. An ultrasound registration was also conducted to observe the depth of the device’s impact in the posterior crural muscles. It was confirmed that deep and substantial tissue disturbances were caused by this impact. According to our findings, myotonometry is an adequate method to establish muscle parameters. Elasticity and stiffness of the *rectus femoris* muscle may determine success in triathlon.

## Introduction

The structural proteins of muscle filaments and the corresponding fascia tissue are contributing to the biomechanical properties of muscles themselves and myofascial structures in general (Grazi, 2008; Palmer et al., 2011; Williams et al., 2012). Kyröläinen et al. (2003) observed the lower-mobility titin band only in the most economical runner in a small group of sprinters. According to Heyward (1998), at age 20–29 the level of superior oxygen consumption among general population in men is >56 ml/kg/min. It has been suggested that VO2max 65 ml/min/kg separates the top-level endurance runners from the rest (Sjödin and Svedenhag, 1985). On the other hand, it would be logical to assume that if the locomotion of an athlete is exceptionally economic due to the beneficial muscle composition and properties, the value of maximal oxygen consumption may not be enormously high.

Economy of locomotion could serve as a predictor of athletic ability, and if this is the case, the biomechanical parameters of muscles must play a role. Muscle properties have been shown to differ among highly and less trained athletes (Fukashiro et al., 2002; Hanson at al., 2012). Stiffness has been found to contribute strongly to the efficacy of the athlete’s performance (Wilson et al., 1994; Kuitunen et al., 2002; Rabita et al., 2008). However, this is not always the case (Slawinski et al., 2008). Muscle stiffness has been reported to differ according to performance ability, and it has been suggested that stiffer muscles allow cyclists to perform better on certain occasions (Watsford et al., 2010). Dumke et al. (2010) demonstrated that muscle stiffness is related to running economy at a speed that approximates endurance competition. Myotonometric research show biomechanical properties correlating to aspects of muscles’ function and competition results (Kim et al., 2016; Hong et al., 2020; Berzosa et al. 2021). Jiménez-Sánchez et al (2018) found, that passive resistive torque of the triceps surae muscle (measured with the isokinetic device) is higher in more stiff, tense and elastic muscle.

During triathlon competitions, running performance seems to be extremely important (Knechtle and Kohler, 2009). Triathletes differ in their parameters from pure endurance runners by larger muscle mass necessary for cycling and swimming (Sleivert and Rowlands, 1996), though. Bonacci et al. (2011a,b) showed that switching from cycling to running is a crucial neuromuscular event that ensures success and singles out the elite from the rest.

Elasticity of specific muscles is defined less in literature. It is a quality describing the ability of the body to resume the initial form after deformation by compression, stretching, twisting or bending. Industrial elastic springs are made from stiff and plastic materials. Cavagna et al. (1971) suggested that elastic energy is stored and released in muscles and tendons during sprints. In wallabies and kangaroos, elastic energy may account for up to half of the performed work at hopping without additional energy cost (Kram and Dawson, 1998). Elasticity is usually mentioned in studies using elastography—detection of share-wave propagation in tissue(s)—, but this propagation depends on the arrangements of the muscular structures (Muthupillai et al., 1995; Gennisson et al., 2005; Eby et al., 2013). In our study a different method and device was used and biomechanical properties like elasticity, stiffness, and the tension in the muscle were established. Ditroilo et al. (2011) showed that elasticity depends on the stretch and that muscles become slacker/more plastic when less stretched. Jarocka et al. (2011) showed that if the contraction force rises gradually, muscle elasticity does not behave accordingly. Instead, it rises rapidly at a low contraction force to a high plateau level while stiffness and tension correlates almost linearly with the generated power. It is important to note that all three parameters are calculated from the same single measurement. According to most recent observations, the elasticity of the muscles in master athletes is lower than in sedentary coevals (Gervazi et al., 2017) and intensive physical exercise significantly increases stiffness in the Achilles tendon (Pozarowszczyk et al., 2017). Most recent work correlations strongly Achilles tendon elasticity (among other parameters) to countermovement jump height (Wdowski et al., 2022).

As the impact from the myoton device is light and given on the skin, there are questions about its ability to reach and describe muscle properties. We quote, “So it is possible that during rest conditions, the Myoton-3 measures mainly the oscillations of the skin and subcutaneous tissue which are provoked by its testing-end.” (Jarocka et al., 2011); “Myotonometry is quick and inexpensive, but tends to be superficial or merely qualitative.” (Eby et al., 2013). On the other hand, “The linear relationship with force output suggested that the device was giving a valid recording of the viscoelastic stiffness of the muscle rather than that of the subcutaneous tissue.” (Bizzini and Mannion, 2003).

This is confirmed by Zinder and Padua (2011), and using newer myoton device by Kelly et al. (2018), and Li et al., (2022). A reliability study made by Bravo-Sánchez et al. (2021) revealed that the thicknesses of all tissues beneath the testing-end (muscle tissue, connective and superficial tissue, adipose tissue) correlated separately with the myoton registered stiffness values. Fröhlich-Zwahlen et al. (2014) obtained analogues results previously. As the effect of the device on tissues was not clear, we decided to study this using an ultrasound (US) registration that allows to detect deep tissue disturbances (Gennisson et al., 2005). This experiment was conducted to answer a single question— does the 0.4 N mechanical impact from the device reach the muscle tissue—as it has not been studied so far.

## Methods

### Atheletes

The study protocol was approved by The Ethics Review Committee on Human Research of the University of Tartu (No. 94/17; 21.05.2001 for athletes and 242/T-21 for the US study). All participants gave their informed consent in writing. Fourteen male triathletes were observed during a period of 36 months. Athletes of the national junior team were studied from whom most had been trained regularly as triathletes under the same coach. Elder triathletes were high rank hobby athletes. The criterion was set that the athlete must have been highly active more than 3 years to be chosen into the study. Their height was from 176 to 192 cm, and their body mass varied from 66.1 to 81.6 kg, age varied between 17–41 years at the beginning of the study. Average values of height, weight and BMI were used.

During the study period, the athletes were tested on VO2max up to 4 times. According to their average VO2max results, the group was divided into half.

Separately, three non-athletes but athletic men with BMIs 27, 26, and 23, were used for the US study.

### Myotonometry

The testing-end of the device with the effective weight P, is placed on the skin surface transversally to the muscle (level S as dotted line; Figure 1 B) compressing the tissue in terms of _Δ_S. The connected to testing-end electromagnetic driver (Figure 1 A) is fired (time-point t_1_) producing a short impulse (t_k_=15ms=t_1_-t_2_) terminating with a quick release at moment t_2_. This generates a mechanical force of 0.4 N (Newton). The device then monitors evoked primary oscillatory waves, as the testing-end stays in contact with tissue surface. Acceleration of the testing-end is recorded at a 3 KHz frequency.

**Figure 1.**
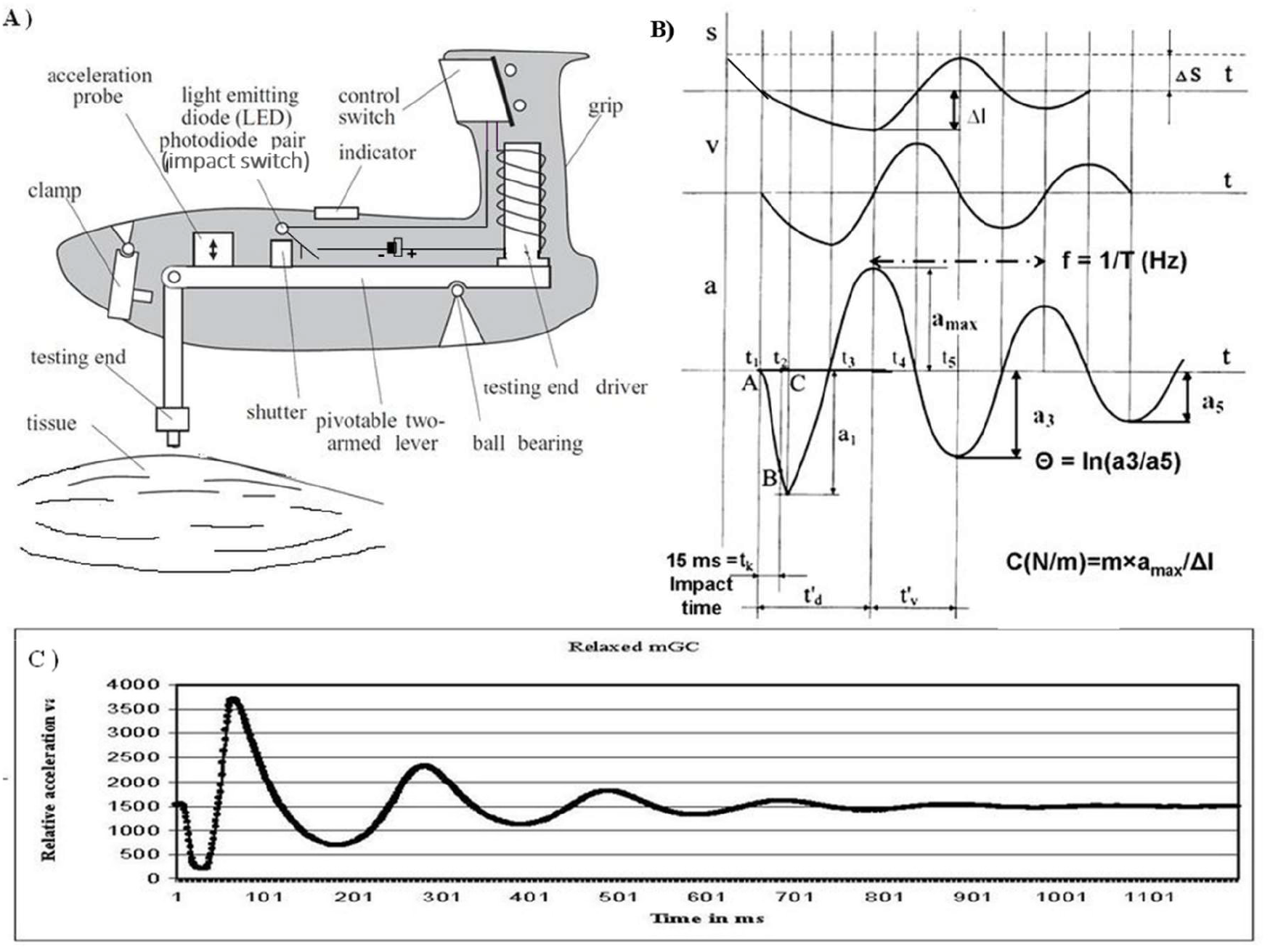
Myotonometry [with courtesy to *Dr* (*Habil. Biol.*) PhD Arved Vain]. A) Working principle of Myoton models 2 and 3. B) Schematic graphs and formulas (see the text). Waveforms of displacement (s), velocity (v), and acceleration (a). C) Graphic presentation of the acceleration values recorded during the measurement.

Elasticity is the ability of a tissue to restore its initial shape and is characterized by the logarithmic decrement of the damped oscillations (Equation 1). The smaller the value, the more elastic is the tissue showing that less energy is lost in each following oscillation. Stiffness reflects the resistance of the tissue to the force that changes its shape (Equation 2). The equipment detects it with the testing-end during the initial impact. The higher the value, the stiffer is the tissue, showing that more energy is needed to modify muscle shape. The frequency of the damped oscillations (Equation 3) characterizes the state of the tissue – the higher the value, the more tense is the tissue. We used Myoton-2 on athletes and Myoton-3 in the US study. The working principle and build-up of both models is the same (Figure 1 A).

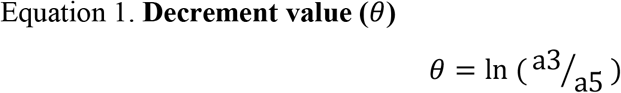

a–amplitude (Figure 1 B). Smaller decrement represent better elasticity, large values higher plasticity

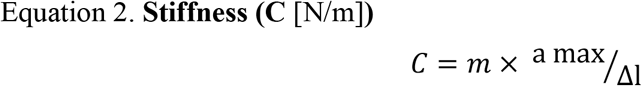

Characterizes the deformation of the muscle caused by the testing-end (Figure 1 B).

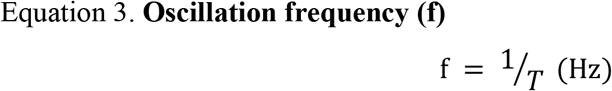

T is the period between the peaks of amax and a4 in seconds (Figure 1 B)

### Procedures

Muscles: BB – *biceps brachii (caput longum)*; TB–*triceps brachii (caput longum)*; BF–*biceps femoris (caput longum)*; RF–*rectus femoris*; TA–*tibialis anterior*; GC–*gastrocnemius (caput mediale)*; LD–*latissimus dorsi*; PM–*pectoralis major (pars sternocostalis)* were measured bilaterally in both relaxed and contracted state while the subjects rested supine or prone, depending on the muscles measured, on a portable massage table. All the measurements were carried out by the same person. The measuring point was marked on the skin at the most prominent point of the muscle belly at contraction (Gavronski et al., 2007). Contraction was standardized simply by the same position of the limb and additional weight (a 2.3 kg dumbbell) was used when the muscles of the upper body and the brachium were measured. A heavier weight caused muscle tremor which disrupted the measurements. To evoke contraction in the muscles of the brachium (BB and TB), the subject was holding his arm at the level of the shoulder and raised it to an angle of 45° from the horizontal axis (measured with a fixed angle), holding a dumbbell. To contract the PM muscle (supine position), the subject stretched out his arm horizontally on the side over the table edge, holding a dumbbell in his hand. For LD (prone position), the subject stretched his arm similarly over his head. The contraction of the BF and the TF muscles was evoked by raising the leg to an angle of 45°. To contract the GC and the TA, the maximum tarsal dorsiflexion and plantarflexion were performed against the fixed table. Most measurements were taken weekly on Wednesday mornings before the training sessions. There were few periods when measurements were taken daily for 3–5 days, also in a field situation before the warm-up for the competition and immediately after the competition. In two longer periods from summer to winter (about five months), no measurements were taken.

The subjects were tested for VO2max up to four times during this period. Standardized complex laboratory tests of functional aerobic capability were performed on a Technogym Runrace HC 1400 (Italy) treadmill. The same gradually increasing load regime was used. After 5 min warm up at speed 8km/h, the starting speed was set to 10km/h and then raised by 2km/h after every 3 minutes up to 14 km/h. From that point, the speed was raised in the same rhythm by 1 km/h until exhaustion. Respiration parameters with O2 and CO2 fractions in the exhaled air were measured using a True Max 2400 (Parvo Medics, USA) computerized complex. The criteria used to confirm that VO2max had been reached was the achievement of the plateau level in O2 uptake with increase in work-rate at the R value greater than 1,1. The group was divided into half according to average oxygen consumption, seven subjects in both groups. Those in the higher group were marked as ‘High’ and the others as ‘Low’.

### Ultrasound Measurement

We made US recordings on the *Musculus gastrocnemius* (GC) in the same direction as the mechanical impact from myoton. This muscle was chosen because it is easily accessible. Due to the unavailability of the same device used on athletes and corresponding Myoton-2 models, we used similar but newer Myoton-3 with same build-up and impact force, for that purpose (Figure 1 A; B). The use of Myoton-3 was also reasoned by the fact that prior to MyotonPRO, this model was mainly used.

An ultrasonic apparatus General Electric LOGIQ E9 with an electronic linear array probe 11L (11 MHz wave frequency with 45 mm scanning length) was used to obtain images of the lateral GC muscle area, which were recorded to a video file at 25 Hz. We placed the US inducer on the ventral surface of the GC along the longitudinal vector of the leg on the belly of the muscle and applied impact along the same line with the US beam. The location of the test-end aside the US probe is marked with an arrow in Figure 2. The location of the measuring point was chosen as homogenous as possible, in the area of the lowest pulsation from vessels registered previously by Color-Doppler.

**Figure 2.**
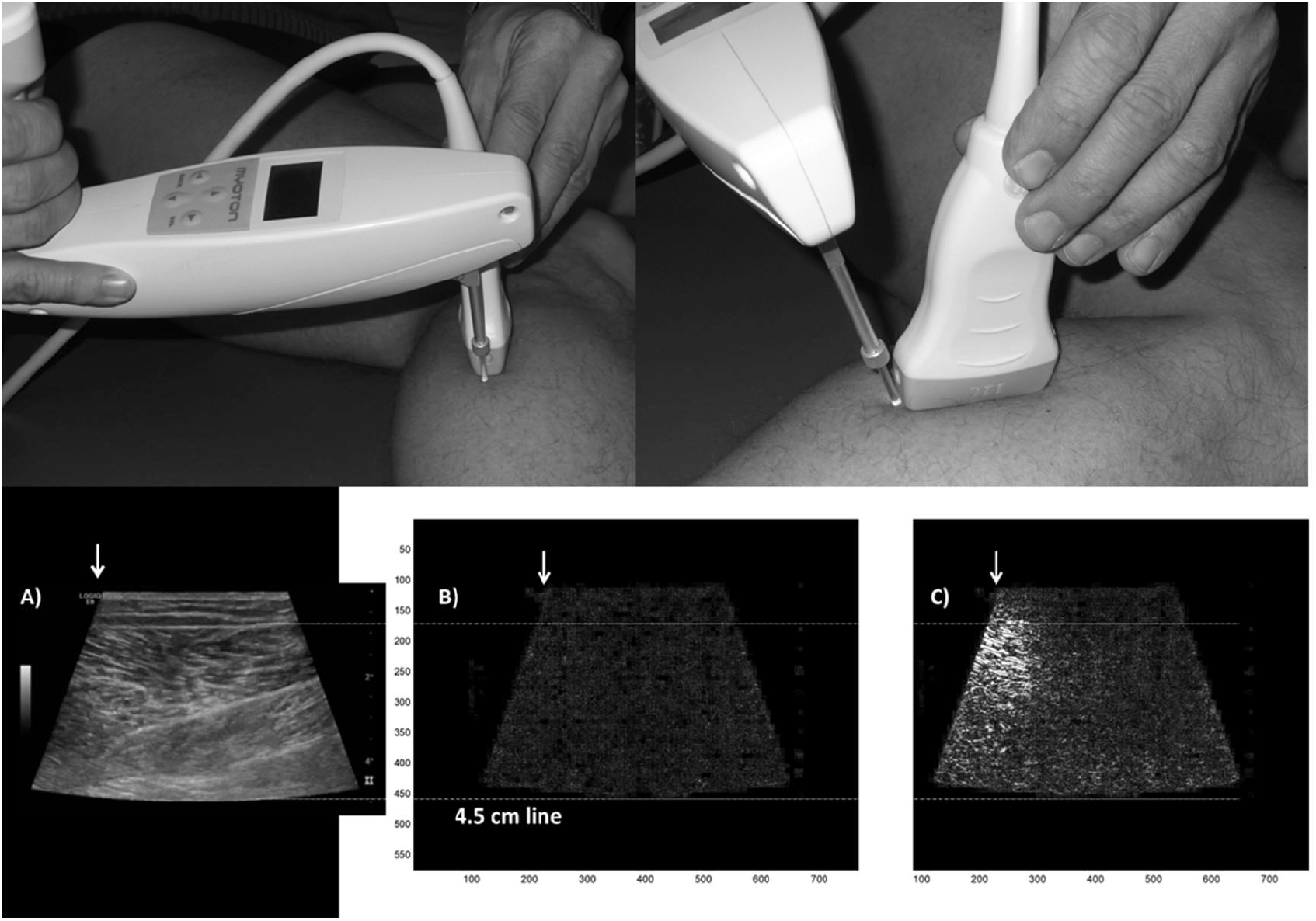
Illustrative placement of devices to receive the ultrasound recording. Tissue reaction on impact. A) An original image from US video; B) Results of image analysis with MatLab before the impact; C) immediate effect of the impact. White arrow–impact site. Brighter white color on image B presents stronger disturbances. Upper guideline marks the superficial aponeurosis of the gastrocnemius muscle, lower guideline marks the depth of 4.5 cm in the tissues

### Image Analysis

To confirm the visual findings, we used Matlab 5.3 software and analyzed recorded video material separating frames from the black and white video clips obtained from the US recording. Extracted frames were divided into pixels on X (768 dots per image) / Y (576 dpi) scale and each dot was monitored on four sequential images establishing the standard deviation of the change on the dark/light scale. New images were generated visualizing the deviations of each monitored dot and higher deviation is presented with the brighter white color (Figure 2).

### Statistics

The Statistica software (version 8; StatSoft Ltd, Bedford, UK) was used. ANOVA was used to compare the general parameters like height, weight, BMI, age and VO2max. The values of the biomechanical parameters in relaxed and contracted states were analyzed separately. According values were pooled either for one person to receive the average “general” value of each biomechanical parameter, or for each muscle. These average values were correlated with VO2max. To compare the muscles between both groups, both sided muscles were considered separately. Non-parametric Mann-Whitney U-test (MW) was used to compare the biomechanical parameters and Spearman Correlation R was also calculated.

## Results

### General Parameters

The age of the athletes did not differ between the groups (Table 1 shows the age at the end of the study; ANOVA F(1,12)=0.13; p=0.728). There was a significant difference in the height and VO2max values between both groups (Table 1). The general biomechanical parameters of muscles did not differ between the groups. The VO2max did not in correlate with body mass, BMI, age, or height.

**Table 1.** General parameters.

| Group | Height (cm) | Body mass (kg) | BMI | Age (years) | O2 ml/kg/min |
| --- | --- | --- | --- | --- | --- |
| High | 188.7±3.8* | 78.8±11.3 | 22.0±2.5 | 24.4±5.4 | 66.4±3.1** |
| Low | 182.0±6.7 | 75.5±7.7 | 22.7±1.8 | 25.9±9.1 | 59.5±2.9 |

*Mean±SD are given. N=7 for each group. Age is given at the study's end year. ANOVA was used and \* $p<0.05$ , \*\* $p<0.0001$ compared to the Low VO2max group are presented*

### Image Analysis from the Ultrasound Recordings

The answer to our study question was that the 0.4 N impact from the myoton device is sufficient to reach the muscles through the skin and subcutaneous tissue causing substantial tissue disturbances in muscles. Representative image is presented as Figure 2C.

### Elasticity

Elasticity improves as the decrement value lowers, hence, if the other parameter grows, the negative correlation reflects better elasticity and if both parameters grow, positive correlation reflects lesser elasticity or higher plasticity in turn. For this analysis we calculated single general value for each athlete (N=14). General decrement value in a relaxed state showed higher plasticity at higher body weight and BMI (weight R=0.73, p=0.003; BMI R=0.86, p<0.0001). Oxygen consumption did not correlate with general elasticity while did with the elasticity of the contracted RF muscle (R=−0.58; p=0.029).

To compare between the High and the Low groups, each muscle was considered separately (N=2×7=14 muscles in both groups). A significant difference was revealed in the average elasticity of the RF in both relaxed (MW test; p=0.035) and contracted (p<0.001) state (Figure 3; Table 2).

**Figure 3.**
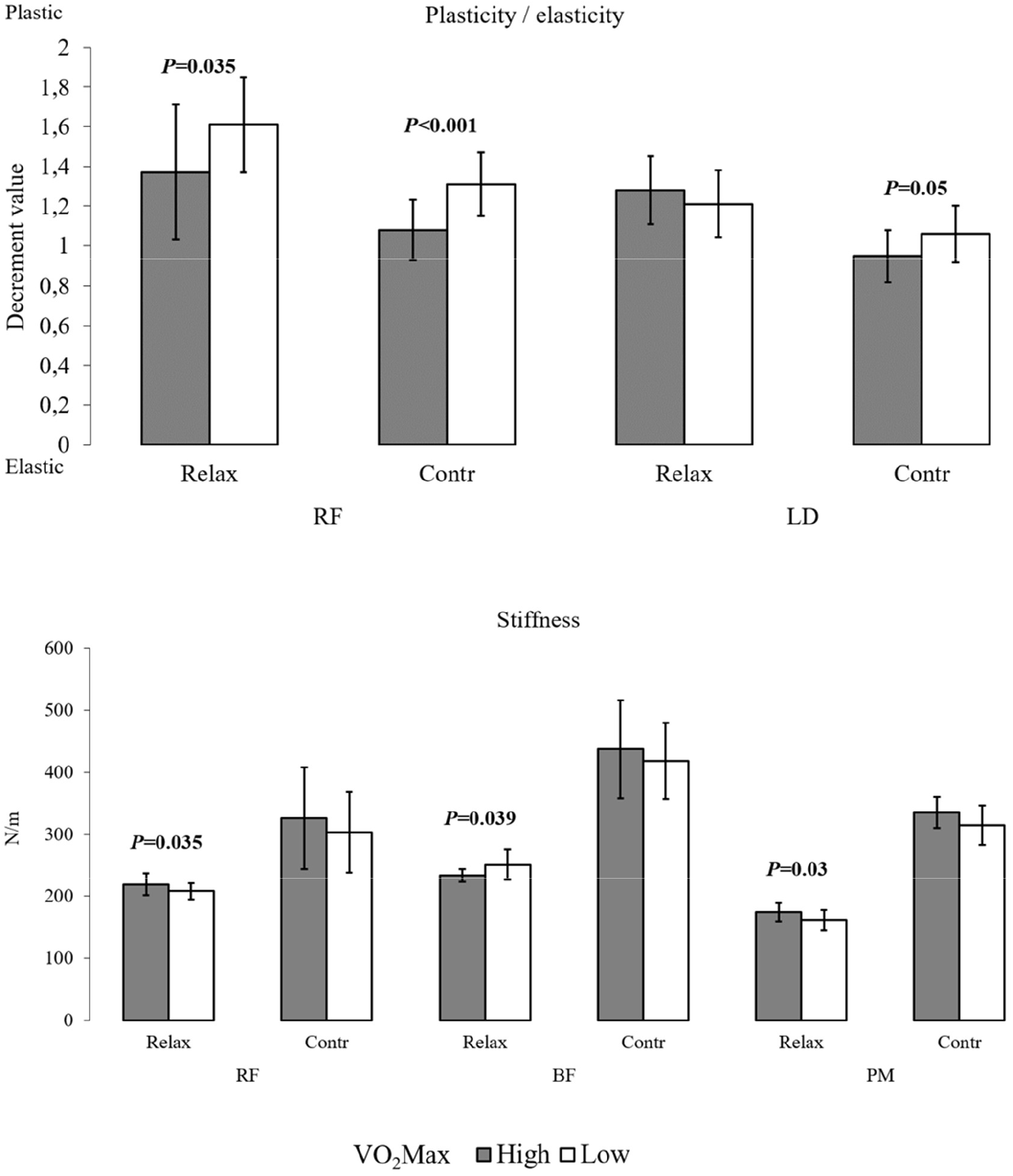
Elasticity and stiffness in significantly different muscles. Differences in muscle biomechanical parameters according to VO_2_Max. Exact data is presented in table 2.

**Table 2.** Biomechanical parameters of skeletal muscles.

| Average ±SD | | Decr. | | Stiff (N/m) | | Frequency (Hz) | |
| --- | --- | --- | --- | --- | --- | --- | --- |
| Mus. | Level | Relaxed | Contracted | Relaxed | Contracted | Relaxed | Contracted |
| BB | High | 1.16±0.16 | 0.85±0.04 | 154.4±11.0 | 218.7±25.4 | 9.30±0.39 | 12.19±0.92 |
| BB | Low | 1.21±0.17 | 0.91±0.13 | 154.9±15.5 | 218.9±32.7 | 9.52±0.60 | 12.28±1.08 |
| TB | High | 1.30±0.13 | 0.94±0.16 | 201.7±15.5 | 307.5±70.3 | 11.65±0.61 | 15.28±1.89 |
| TB | Low | 1.33±0.16 | 0.95±0.09 | 191.0±26.1 | 322.6±60.6 | 11.34±1.15 | 15.77±1.84 |
| LD | High | 1.28±0.17 | 0.95±0.13* | 154.5±11.3 | 381.3±71.2 | 8.74±0.47 | 17.24±2.12 |
| LD | Low | 1.21±0.17 | 1.06±0.14 | 154.7±15.3 | 350.2±42.4 | 8.98±0.44 | 16.80±1.23 |
| PM | High | 1.37±0.25 | 0.75±0.10 | 174.2±15.1* | 334.8±24.8 | 10.29±0.55 | 15.73±0.57 |
| PM | Low | 1.32±0.28 | 0.89±0.23 | 161.2±16.4 | 314.7±32.0 | 9.88±0.55 | 15.19±0.98 |
| BF | High | 1.50±0.45 | 1.24±0.14 | 233.4±10.5* | 436.8±78.6 | 13.33±0.54 | 18.72±2.23 |
| BF | Low | 1.53±0.17 | 1.25±0.24 | 250.8±24.4 | 418.3±61.7 | 13.88±1.14 | 18.22±1.55 |
| RF | High | 1.37±0.34* | 1.08±0.15* | 219.2±17.7* | 325.5±81.7 | 11.98±0.84 | 15.93±2.44 |
| RF | Low | 1.61±0.24 | 1.31±0.16 | 207.9±13.2 | 302.7±65.0 | 11.40±0.63 | 15.23±2.23 |
| TA | High | 0.89±0.11 | 0.78±0.09 | 399.7±42.8 | 822.4±100.7 | 18.52±1.42 | 26.82±2.16 |
| TA | Low | 0.84±0.11 | 0.81±0.11 | 383.7±50.9 | 835.7±112.8 | 18.57±1.57 | 27.62±2.14 |
| GC | High | 1.05±0.17 | 1.02±0.16 | 215.8±28.0 | 366.6±69.0 | 11.33±1.38 | 17.07±1.60 |
| GC | Low | 1.13±0.20 | 1.06±0.17 | 224.2±31.5 | 370.4±52.4 | 11.74±1.18 | 17.43±1.58 |

*Average values are presented with ± SD; N=14 (7 persons x 2 muscles) for each group.*
*The Mann-Whitney U-Test was used. \* means a significant difference ( $p < 0.05$ ) with the*
*respective Low group parameter*

In a relaxed state, the RF muscle was significantly more elastic in High group on both sides (left side: High 1.16±0.13 v. Low 1.78±0.20, p=0.002, N=7; right side: High 1.31±0.18 *v*. Low 1.71±0.22; p=0.002; N=7). No difference between the groups was observed for contraction (left side: High 1.07±0.2 *v*. Low 1.28±0.18, p=0.06, N=7; right side: High 1.14±0.17 *v*. Low 1.27±0.16, p=0.18, N=7).

Additionally, the LD muscle in a contracted state showed a significant difference between the groups (p=0.043; Figure 3; Table 2), but showed no correlation with oxygen consumption.

### Stiffness and Frequency

Both general parameters correlated well with body height (Height/Freq R=0.68, p=0.008; Height/Stiffness R=0.72, p=0.004; N=14) but no difference was evident when taller High VO2max group was compared to the shorter Low group.

Between the two groups, three muscles showed a significant difference in relaxed state and only in stiffness—both thigh muscles (BF p=0.039 and RF p=0.035) and PM (p=0.03; Table 2). Interestingly, the antagonistic thigh muscles showed the opposite behavior—RF was stiffer in the High group and BF in the Low group.

## Discussion and Conclusions

To our knowledge, this is the first study on the actual tissue impact of myotonometry in general and Myoton-3 device in particular. We stress that the aim of the US experiment was simply to assess the adequacy of a 0.4 N mechanical impact to reach the muscles. As a result, we observed deep tissue disturbances as the impact penetrated the skin and subcutaneous tissues in a relaxed state of the muscle. Our conclusion was supported by the finding that these strong disturbances were observable on 11 consecutive newly generated images that makes the duration of the impact ≈ 440 ms (1000 ms / 25 frames per second × 11 frames), which is consistent with the approximate duration of strong oscillations (Figure 1 C). It was not our aim in this study to match the results from other measurement with the findings on video-recordings, neither to make any quantitative analysis nor find any correlations in this matter. These findings are also in same line with our jet unpublished data. According to our findings we can firmly state, that the device is appropriate for describing specifically muscle properties. This was suggested already by Bizzini and Mannion (2003). It is our suggestion that this is possible due to the incompressible nature of soft-tissues consisting mainly of water (Grazi, 2008) allowing equalizing the pressure for Korhonen et al. (2005) have proven the compartmental pressure to correlate well with regional biomechanical parameters.

Our second main finding was that oxygen consumption correlates with the biomechanical properties of the RF muscle, allowing describing up to 1/3 of the general VO2max with the elasticity of this muscle in studied triathletes. This is supported by the difference in elasticity of this muscle between the High and Low groups and in addition, the analysis between general decrement values from activated muscles with VO2Max showed R=−0.45 with p=0.1. The main reason for the prominence of RF muscle is clearly the cycling and running events. Hein and Vain (1998) showed that motion over the hip joint is related to the biomechanical properties of the agonist-antagonist muscles. They stated that the knee extension range of motion was primarily related to the elasticity of contracted posterior group of thigh muscles and that the tension (oscillation frequency) of *m. semitendinosus* was most responsible for limiting trunk forward flexion. This seems to support our finding that BF is stiffer in low endurance capable group, reflecting impeded locomotion over hip and/or knee joints when cycling or running. Zajac (2002) explains, that the muscles producing highest work output during pedaling are uniarticular hip extensors (e.g. *gluteus maximus*) with anterior thigh muscles i.e. the uniarticular knee extensors (*vastus medialis, lateralis*, and *intermedius*) and that these muscles deliver their work output to both the crank and leg during leg extension. It has been shown that in cycling, the leg muscles exploit different metabolic paths than the ones of arms and that legs are responsible for using most of oxygen (Volianitis and Secher, 2002; Calbet et al., 2007). This knowledge seems to support our findings here. Cycling has been suggested to be crucial for successful running in triathlon (Bonacci et al., 2011a,b). Still, Hausswirth et al., (1999) pointed out that a specific tactical decision, like drafting etc., allows triathletes to save considerable energy for running, suggesting also that this could mainly benefit good runners. Chapman et al. (2008) and Candotti et al. (2009) suggested that triathletes have a more robust cycling technique and that their recruitment of the BF muscle differs from cyclists. In transition from cycling to running, the recruitment of the *tibialis anterior* muscle seems not to be affected (Eby et al., 2013) and we did not see this muscle (neither the GC) to emerge. A recent study shows that in cyclists sprinting event, anterior thigh muscle plasticity accumulatively raised in a course of competition and measured leg flexors showed greater decrement values (lower elasticity/higher plasticity) than extensors. The 200m flying sprint—similarly to continuous maximum effort, affected most the elasticity of anterior thigh (Klich et al., 2020; 2022). Differently from cyclists or triathletes, in endurance runners the VO2max is lower if it is measured on veloergometer compared to treadmill (Kohrt et al., 1989; Hue et al., 2000; Basset and Boulay, 2003; Davis et al., 2006; Millet et al., 2009; Carey et al., 2009). Therefore, it seems that running only does not allow fulfilling the cycling task with the same efficacy, while triathletes are able to run and cycle with similar intensity. In future, comparing triathletes with cyclists and runners may provide more knowledge about these issues.

Tissue elasticity (in a relaxed state) was strongly associated with both body weight and BMI in the way that more body mass was accompanied with higher plasticity. Negative correlation between the RF and BF muscle thickness (established by US) and decrement value was observed by Fröhlich-Zwahlen et al. (2014) although they misinterpreted the results concerning the decrement value related to elasticity—<u>the higher the decrement value, the less elastic or more plastic is the tissue</u>. This means, that they observed higher elasticity at higher thickness in thigh muscles in non-athletes.

From previous research, it can be concluded that if a muscle is contracted, the force can be derived at least to some extent correctly from the stiffness or tension if measured alone (but not from elasticity; Jarocka et al., 2011). As these two parameters also depend on the stretch (Ditroilo et al., 2011; Huang et al., 2018), the correlations between biomechanical parameters and the height allow us to suggest that in taller people muscles may be naturally more stretched and that fore stiffer. This may be the reflection of the effect of Gravity force on standing posture.

We also observed a trend between the VO2max and the elasticity in contracted BB (R= −0.47; p=0.09). Accompanied with the LD muscle showing better elasticity in a contracted state in the High group, these findings in these two muscles may reflect the effect of swimming trainings. On the other hand, as the contraction in our study was called out with a simple task, the excessive performance in the High group may have interfered causing these differences, for both muscles showed improved elasticity accompanied by slightly higher stiffness and tension (Jarocka et al., 2011). It is a shortcoming of our work that the relaxed and contracted states were not registered by any other objective measures like EMG or generated force. It was our decision in order to simplify the procedure. As we calculated average over all the measurements, each single one less affects received values. Most recent study by Mencel et al. (2021) cleared the doubts about accuracy of measurement point. According to their findings, the deviation up to 20% distally from muscle-belly is irrelevant in large muscles. Mencel et al. used MyotonPRO, still the general principle of oscillatory measuring of the mentioned parameters is the same in both devices.

There are some aspects in our study that we want to emphasize. First, the study group is small. This may compromise the ability to translate our main finding to other sports. Important aspect is that all values are pooled over all three years. In each athlete, more than 30 measurement sessions were conducted. During this period, we observed that muscle parameters are strongly affected by a single training or competition event (data not published). Pooling of data abolishes these scattering effects. It is important to note that testing muscles only in a relaxed state definitely does not reveal all the valuable information and we recommend measuring also in the activated state. Contraction must be sufficient and standardized for each muscle, though.

Despite the rapidly growing number of publications on myotonometry, available material is scarce. Only a few aspects of these properties are covered in the literature, most of them describing stiffness or tension as properties and measurements made from tendons. Elasticity seems to be more stable and less volatile biomechanical property (Vain et al. 2015, Julià-Sánchez et al. 2020), that is also our experience, and authors are not reporting the results. That fore there is an urgent need for more studies using this methodology. According to most recent publication by Wdowski et al. (2022), higher countermovement jump performance is enabled by higher elasticity (smaller decrement value), stiffness, and tone, with smaller creep-ability and quicker stress-relaxation times measured from Achilleus tendon. Two latter parameters are unique to newest myotonometric device.

In conclusion, myotonometry is a simple method to obtain objective information about the state and condition of muscles with the potential to monitor the training process. This would be beneficial in optimizing trainings, detecting muscle soreness in order to avoiding traumas, and monitoring rehabilitation. Large groups of athletes, separating sprinters and endurance athletes, in uniform events as running, cycling, and swimming are good candidates for future research.

## Acknowledgements

The authors are greatful to Arved Vain, PhD, Dr. Habil. Biol., for his expertise; Prof. Vahur Ööpik, Dr Jüri Laasik, Kalle Karelson, Cand. of Biology, and Saima Timpmann, MSc, for their consultations and help. Thanks go to Jüri Käen, the coach of the triathletes; Inge Ringmets, MSc, and PhD student Kristiina Rajaleid for their help with the statistics. Also to MD Adu Simisker for his expertise in ultrasonography.

## Notes

**CONFLICT OF INTEREST**, a. There is no duplicate publication elsewhere of any part of this work, there is no closely related paper published by us previously in this or in other journals. b. There is no conflict of interest of any sort between the authors and the study presented. This study was not funded by the interested parties, or by any other foundation. c. All authors were fully involved in the study or preparation of the manuscript. The typescript has been read and agreed by all authors. d. There are no competing interests bound to this study.

### Competing Interest Statement

The authors have declared no competing interest.

